# The neural basis of hand choice: An fMRI investigation of the Posterior Parietal Interhemispheric Competition model

**DOI:** 10.1101/409565

**Authors:** Aoife M. Fitzpatrick, Neil M. Dundon, Kenneth F. Valyear

## Abstract

The current study investigates a new neurobiological model of human hand choice: The Posterior Parietal Interhemispheric Competition (PPIC) model. The model specifies that neural populations in bilateral posterior intraparietal and superior parietal cortex (pIP-SPC) encode actions in hand-specific terms, and compete for selection across and within hemispheres. Actions with both hands are encoded bilaterally, but the contralateral hand is overrepresented. We use a novel fMRI paradigm to test the PPIC model. Participants reach to visible targets while in the scanner, and conditions involving free choice of which hand to use (Choice) are compared with when hand-use is instructed. Consistent with the PPIC model, bilateral pIP-SPC is preferentially responsive for the Choice condition, and for actions made with the contralateral hand. In the right pIP-SPC, these effects include anterior intraparietal and superior parieto-occipital cortex. Left dorsal premotor cortex, and an area in the right lateral occipitotemporal cortex show the same response pattern, while the left inferior parietal lobule is preferentially responsive for the Choice condition and when using the ipsilateral hand. Behaviourally, hand choice is biased by target location – for targets near the left/right edges of the display, the hand in ipsilateral hemispace is favoured. Moreover, consistent with a competitive process, response times are prolonged for choices to more ambiguous targets, where hand choice is relatively unbiased, and fMRI responses in bilateral pIP-SPC parallel this pattern. Our data provide support for the PPIC model, and reveal a selective network of brain areas involved in free hand choice, including bilateral posterior parietal cortex, left-lateralized inferior parietal and dorsal premotor cortices, and the right lateral occipitotemporal cortex.

## 1. Introduction

Deciding which hand to use to perform actions is one of the most fundamental choices humans make, and yet the brain mechanisms that mediate hand choice are poorly understood. According to traditional accounts of decision-making, the brain systems governing choices are separate from those that are responsible for the sensory guidance and control of actions (Tversky and Kahneman, 1981; Padoa-Schioppa and Assad, 2006). Accumulating data from multiple domains challenge this view, however, at least with respect to those decisions that determine actions, and suggest that those brain areas important for the control of actions also contribute to action choices (Cisek and Kalaska, 2010; Christopoulos et al., 2015a).

Convergent evidence implicates areas within the posterior parietal cortex (PPC), and interconnected premotor areas, as critical for the planning and control of actions (Kalaska et al., 1997; Wise et al., 1997; Culham and Valyear, 2006). These parietofrontal circuits are responsible for transforming sensory information to motor parameters for the control of actions (Jeannerod et al., 1995; Rizzolatti and Luppino, 2001). This information is available in the neural response patterns within these areas before movements are initiated, and later within primary motor cortex, consistent with their necessary role in action planning and control (Crammond and Kalaska, 1996; Umilta et al., 2007; Schaffelhofer and Scherberger, 2016).

More recently, it has been suggested that these same parietofrontal areas causally contribute to action selection. The very same neural populations responsible for specifying the sensorimotor parameters necessary for the control of actions appear to mediate action choices (Cisek and Kalaska, 2005; Hanks et al., 2006; Scherberger and Andersen, 2007; Pesaran et al., 2008; Pastor-Bernier and Cisek, 2011; Thura and Cisek, 2014; Christopoulos et al., 2015b). These data form the bases of the Affordance Competition Hypothesis (Cisek, 2006, 2007; Cisek and Kalaska, 2010). According to this model, action choices are made by resolving competition between concurrently activated neural populations within parietofrontal areas that specify the spatiotemporal parameters of possible actions.

Motivated by the Affordance Competition Hypothesis, and on the basis of our recent fMRI evidence (Valyear and Frey, 2015), we propose a new systems-level model of human hand selection: The Posterior Parietal Interhemispheric Competition (PPIC) model (Figure 1). Our recent fMRI data suggest that specific areas within bilateral posterior intraparietal and superior parietal cortex (pIP-SPC) represent actions in hand-specific coordinates, and are predominantly contralaterally organized (Valyear and Frey, 2015). These response properties – hand-specific encoding and graded contralateral organization –, together with the population-level neural response principles defined by the Affordance Competition Hypothesis (Cisek, 2006), constitute the essential constraints of the PPIC model.

**Figure 1.**
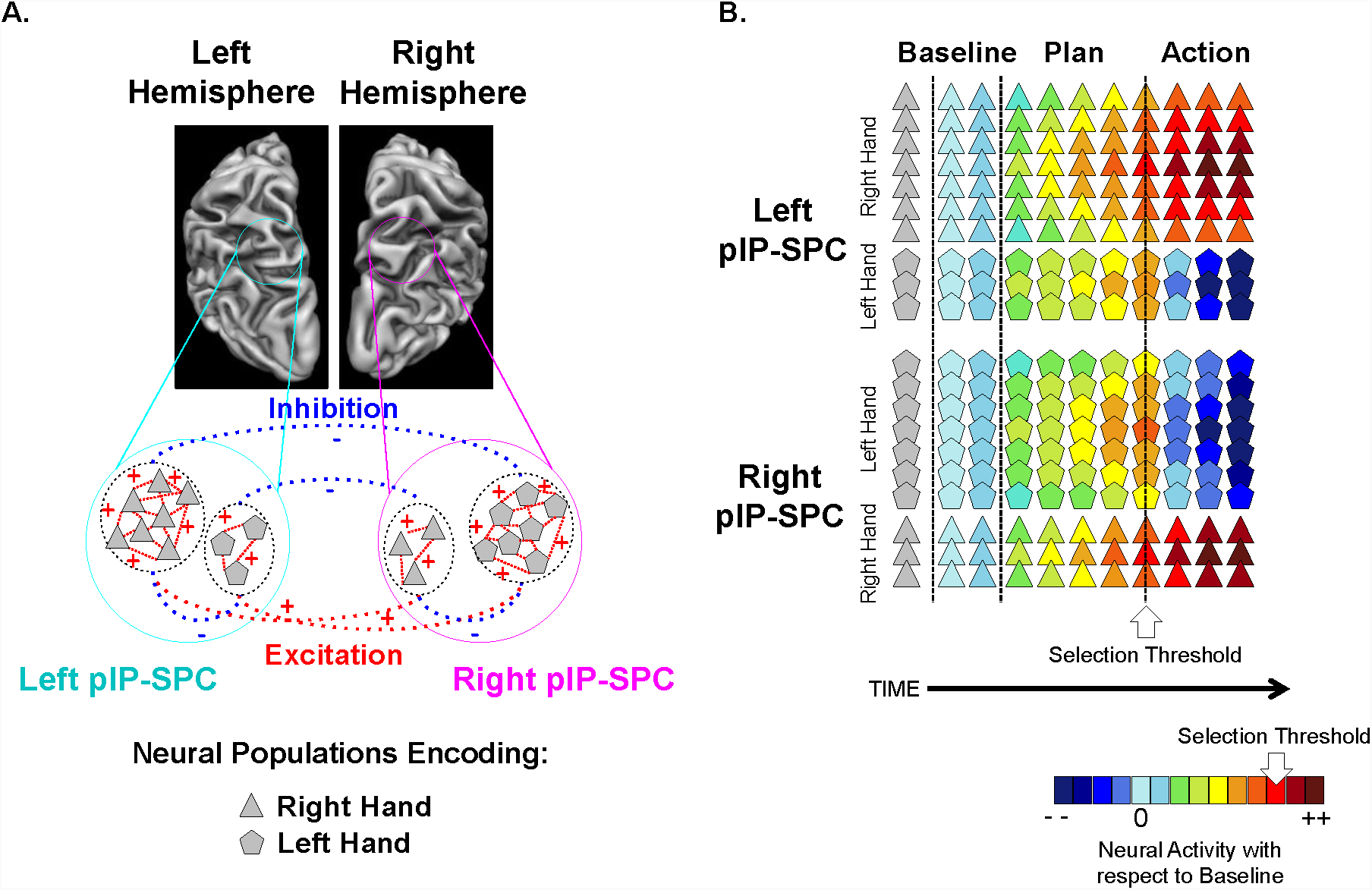
The PPIC model of hand selection. **(A)** Neural populations within pIP-SPC encode actions in hand-specific terms, and a greater number of cells encode actions with the contralateral hand. Cells encoding actions with the same hand excite one another while those that encode actions with the opposite hand inhibit one another. **(B)** Here we show an example of how activity changes in these areas over time in a case where the right hand is selected. During the planning phase the activity of all cell-types increase. The rate of increase depends on various factors, including target location. In this example, those cell populations encoding the right hand show a steeper rate of increase, and reach suprathreshold-activity-levels first. Once threshold is reached, the activity in these cell populations further increases and the spatiotemporal parameters of the actions they encode are selected, while opposing cell populations are robustly inhibited.

Neural populations within pIP-SPC are hypothesized to specify action plans in hand-specific coordinates, and compete for selection across and within hemispheres. Actions with either hand are represented bilaterally, but within each hemisphere a greater proportion of neural populations represents actions with the contralateral hand. Those populations encoding action plans with the same hand excite one another while those that represent actions with the opposite hand inhibit one another. When the activity levels of one population exceed a specific threshold, the parameters of the actions encoded – including the parameter ‘hand’ – are ‘selected’, and competing populations are inhibited.

Distinct from the Affordance Competition Hypothesis, the PPIC model focuses on hand selection, and specifies interhemispheric competition between neural populations encoding hand-specific plans. The predominate contralateral organization of the underpinning neural architecture is an essential feature of the model. This organization drives the proposed interhemispheric competition, and imposes unique constraints on the predictions of the model. Areas within pIP-SPC should not only preferentially respond during conditions involving hand choice, but also for actions made with the contralateral hand.

Findings from a study by Oliveira et al. (2010) provide compelling evidence for the causal involvement of human PPC in hand choice, and suggest an underlying competitive process. Participants used either hand to reach to visual targets presented in left and right hemispace, and the point in target space where the use of either hand was equally probable – the point of subjective equality (PSE) – was estimated. Consistent with a competitive process, response times to initiate actions were prolonged for reaches to targets near the PSE, and these effects were specific to when participants had to choose which hand to use. Further, TMS to the left hemisphere PPC increased the likelihood of reaches made with the left hand. Conversely, TMS to the right PPC did not influence hand choice. The data were interpreted as evidence that hand choice involves resolving competition between lateralized action plans localized within the PPC.

The current study tests the PPIC model, and the hypothesis that bilateral pIP-SPC plays an important role in choosing which hand to use to perform actions. Participants reach to visual targets while lying in the MRI scanner (Figure 2A). In one condition, they are free to choose which hand to use (Choice), while in a second condition hand-use is instructed (Instruct). Targets are arranged symmetrically about the midline of the display, grouped near the centre (Central) and lateral edges (Lateral) of the display.

**Figure 2.**
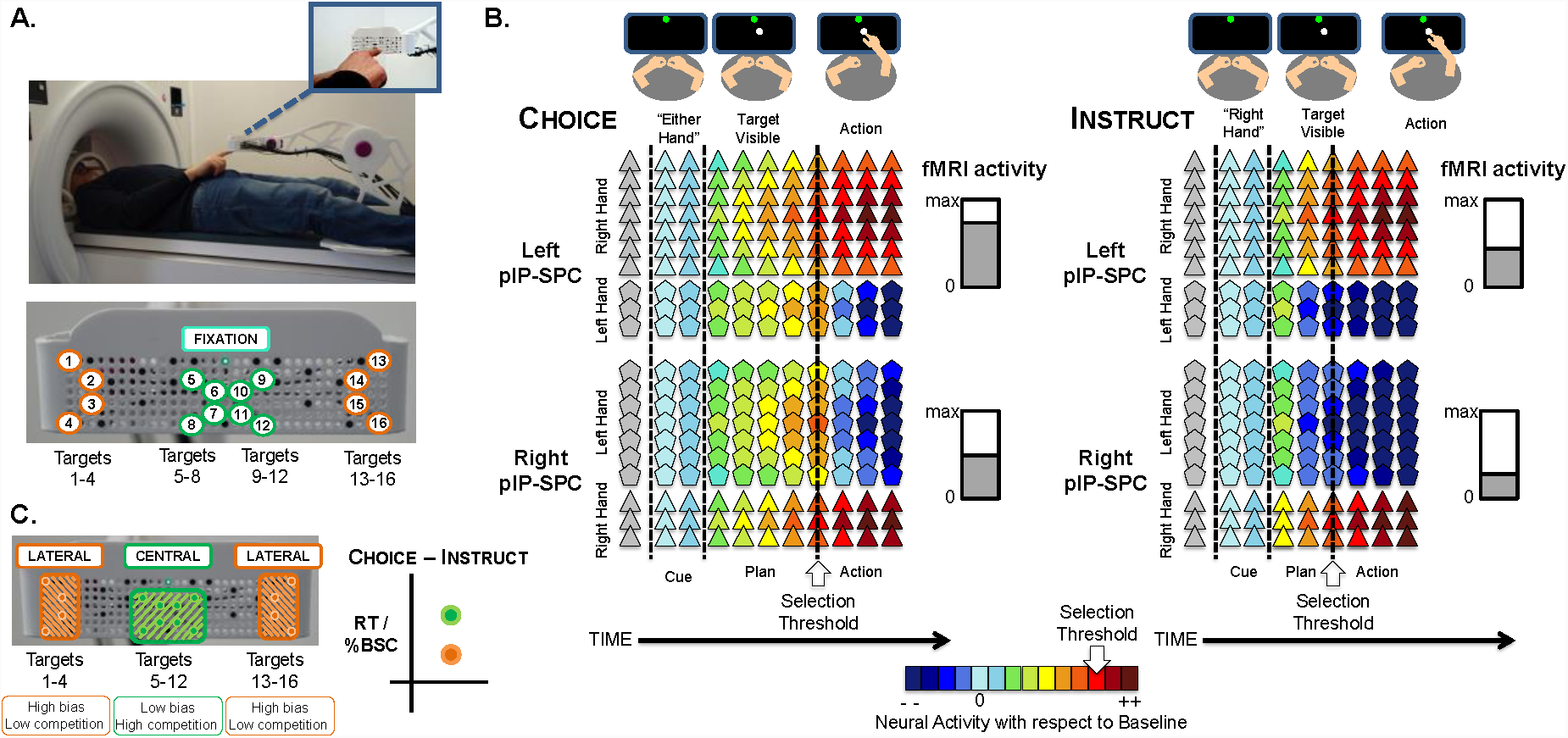
Methods and predictions. **(A)** Optical fibres are fitted to a display module and transmit light to provide 16 targets for reaching, arranged symmetrically around the midline of the display. Targets are presented at left/right Central or Lateral positions within the display. **(B)** The PPIC model predicts a main effect of Task (Choice > Instruct) and a main effect of Hand (Contralateral > Ipsilateral) within bilateral pIP-SPC. Neural populations encode hand-specific action plans, and within each hemisphere, the contralateral hand is overrepresented. Hand choice is determined by resolving competition between active populations. In this example, a Central target is presented and a right-hand response is selected. In the Instruct condition, the competitive process is supervened. This results in reduced fMRI activity levels and RTs relative to the Choice condition. Critically, Choice and Instruct conditions involve the same actions and visual stimuli. **(C)** Hand choice is biased by target location, as a consequence of differing biomechanical costs. Lateral targets represent a high bias, favouring the use of the ipsilateral hand. Stronger bias predicts weaker competition. Central targets represent similar biomechanical constraints for the use of either hand; low bias, and thus high competition. RTs and fMRI activity levels are expected to reflect this gradient: Greater choice-costs (Choice > Instruct) are predicted for Central versus Lateral targets.

The PPIC model makes several specific predictions (Figure 2B/C). First, bilateral pIP-SPC should respond preferentially for the Choice versus Instruct conditions. Critically, in-scanner videos are used to match subject’s behaviour between Choice and Instruct conditions. Differences in activity levels between these conditions are not attributed to visual (or visual-attentional) or motor confounds. Second, bilateral pIP-SPC should respond preferentially for actions made with the contralateral hand – the left hemisphere pIP-SPC should respond more robustly for the selection and use of the right hand, and the right hemisphere pIP-SPC should respond more robustly for the selection and use of the left hand. Third, the anatomical specificity of these effects should correspond with areas previously implicated in the transformation of visual information to motor commands for the control of the arm for reaching.

A final set of predictions is tested. Intermanual differences in biomechanical and energetic consequences, related to the inertial properties of the arm (Gordon et al., 1994), bias both hand (Habagishi et al., 2014; Schweighofer et al., 2015) and arm-movement (Sabes and Jordan, 1997; Cos et al., 2011; Dounskaia et al., 2011) choices. When reaching to targets in either hemispace, the hand that is on the same side of space as the target is favoured, and this bias increases with target laterality (Stins et al., 2001; Oliveira et al., 2010; Valyear et al., 2018). As a consequence, Lateral targets in our display should favour the use of the hand in ipsilateral hemispace, while Central targets should represent more ambiguous choices. This gradient leads to specific predictions within the framework of the PPIC model. Lateral compared with Central targets are predicted to represent more sharply defined reach possibilities, and as a consequence, fewer competing neural populations will be activated and suprathreshold levels will be exceeded sooner – i.e. high-versus low-levels of hand-choice-bias predict low-versus high-levels of competition (Figure 2C). These differences are expected to drive down choice-costs for reaches to Lateral versus Central targets. Both response times (RTs) and fMRI activity-levels are predicted to reflect this pattern: (Choice-Central > Instruct-Central) > (Choice-Lateral > Instruct-Lateral), and these fMRI effects should localize to bilateral pIP-SPC.

## 2. Materials and Methods

### 2.1. Participants

24 individuals participated in the study. One participant’s data was excluded as they reported increasing levels of anxiety and discomfort during scanning, and discontinued testing after four functional runs. The remaining 23 participants (12 female; mean age = 23.2 ± 3.9 years, age range = 20 to 38) were right-handed according to a modified version of the Waterloo Handedness Inventory (Steenhuis and Bryden, 1989; scores range from −30 to +30) (mean score = 23.7 ± 6.2, range = 2 to 30). The experiment took approximately three hours to complete (including pre-scan training), and participants received financial compensation. An additional eight participants completed the pre-scan training (see section 2.4).

All participants were naïve to the goals of the study, and had normal or corrected-to-normal vision, with no history of psychiatric illness. One participant reported prior clinical diagnoses of mild developmental dyspraxia, with no symptomology in adulthood. All participants provided informed consent in accordance with the Bangor University School of Psychology Research Ethics Committee.

### 2.2. Stimuli and presentation setup

Using a custom-built apparatus, targets for reaching were presented to subjects while lying supine in the MRI scanner (Figure 2A). Optical fibres were fitted to the display module of the apparatus (17.5 cm x 6 cm), and used to transmit light to provide 16 targets for reaching, viewed via mirrors mounted to the scanner head coil. Active fibres were symmetrically configured within the display. This organisation ensured that target locations were represented equally across space. Specifically, 8 targets were positioned to the left and right of midline, and within each hemispace, four targets were positioned near the midline (Central), and four targets were positioned near the lateral edges (Lateral) of the display (see Figure 2A). An additional 22 inactive fibres were included, pseudo-randomly arranged, and perceptually identical to the 16 active fibres. This was done to reduce the likelihood that participants would identify and memorize the active target configuration.

The display was adjusted so that all targets were comfortably reachable with either hand with minimal need to move the upper arm or shoulder. Depending on the participant’s arm length, the display distance from the eyes was ∼95 cm. Lateral targets were 7.6 cm (4.6°) and 6.6 cm (4.0°), and Central targets were 1.6 cm (0.97°) and 0.6 cm (0.36°) on either side of the display midline (visual angles are based on a display-to-eye distance of 95 cm, as calculated for one participant). Figure 3A shows target distances from the midline of the display.

**Figure 3.**
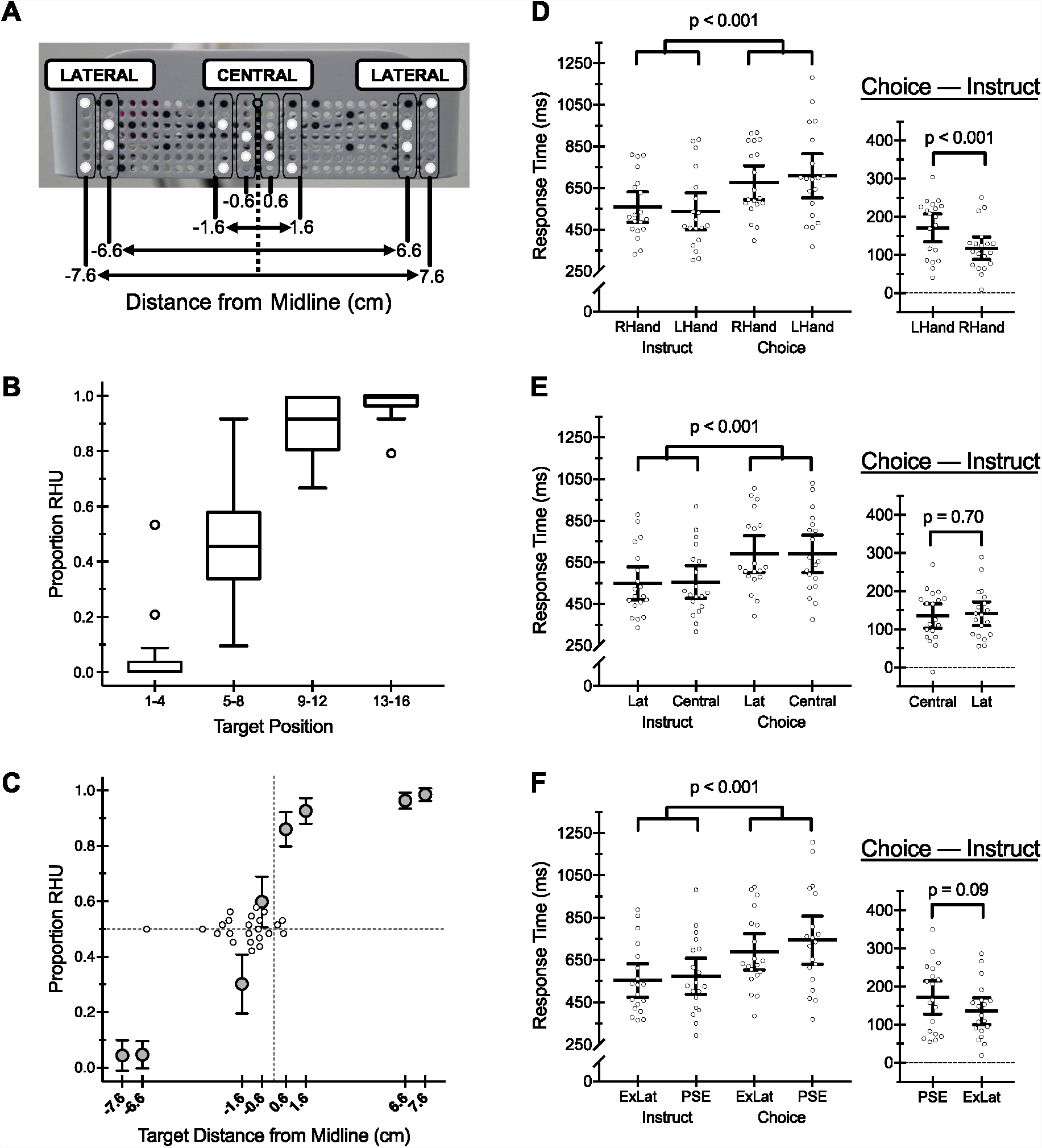
Behavioural results. **(A)** Target space defined as lateral distances from the midline of the display. **(B)** Boxplots showing the proportion of right hand use (RHU) per target quadrant. The lines within boxplots indicate the medians, the upper and lower edges indicate the third and first quartiles, respectively, and the error bars indicate the maximum and minimum data points (excluding suspected outliers). Suspected outliers (1.5*interquartile range above the third quartile or below the first quartile) are shown as unfilled circles. **(C)** Group mean proportions of RHU as lateral distances from the midline. Error bars reflect 95% confidence intervals (CIs). Individual-level PSEs are superimposed on this plot, indicated as unfilled circles. **(D)** Group (N=19) mean RTs as a function of Task and Hand (left), and group mean Choice – Instruct RT differences (right) are shown. Error bars reflect 95% CIs. Individual-level data are shown as unfilled circles. **(E/F)** Same as (D), but showing RTs as a function of Task and Target Location: (E) Central, Lateral; (F) PSE, ExLat.

Participants held down response keys with the index finger of either hand in the rest position. The horizontal midline of the response pad was centred with the horizontal midline of the display module, and secured to the participant’s abdomen near their waistline. In the rest position, the participant’s left and right hands were 3.75 cm lateral to the horizontal midline of the display module. i.e., at rest, central targets were medial to either hand, and lateral targets were lateral to the nearest (ipsilateral) hand. Supplementary Materials include examples of in-scanner videos of participants performing the task.

The apparatus remained outside the scanner bore with the participant localized to the isocenter of the magnetic field. Presentation software (Version 17.2, build 10.08.14) was used for stimulus presentation and behavioural response collection. An MR-compatible infrared-sensitive camera (MRC Systems GmbH) was used to record in-scanner behaviour for offline analyses (see section 2.7.1).

### 2.3. Procedure

At rest, participants fixated a green coloured light-emitting diode (LED) transmitted via an optical fibre positioned in the middle and upper part of the display module (Figure 2A). Trials began with a 600ms duration audio cue: “Left Hand”, “Right Hand”, “Choose”. This was followed by a 200ms delay, and the illumination of a single target. Target illumination lasted for 1200ms. Participants were instructed to reach to targets as soon as they were illuminated, and to fixate targets during reaching. Actions were minimal-amplitude movements, involving mainly the wrist, fingers and thumb, and were approximately 1-3s in duration. Smooth movements, made at comfortable speeds were emphasized. Participants have full-vision available during movements, and thus have visual feedback of their moving limb. Example videos of participants performing the task in the scanner are provided (see Supplemental Materials). Trials were separated by 10s intervals, from target illumination offset.

A slow event-related design was used for two main reasons. First, although perhaps more robust, a block design would be more susceptible to accumulative effects of fMRI-RS (or fMRI-adaptation) due to repeated use of the same hand, and in the case of the Instruct condition, repeated implementation of same rule. This would bias the Instruct condition to have reduced fMRI activity levels (fMRI-RS), and thus make interpretation of our predicted Choice > Instruct effects problematic. Second, a slow event-related design can reveal differences in baseline levels of activity between conditions that may arise prior to trial onsets, and otherwise complicate results interpretation. As such, we were able to rule out the possibility that such differences could account for our data (see Figures 5 and 6, event-related averaged time-course data).

**Figure 4.**
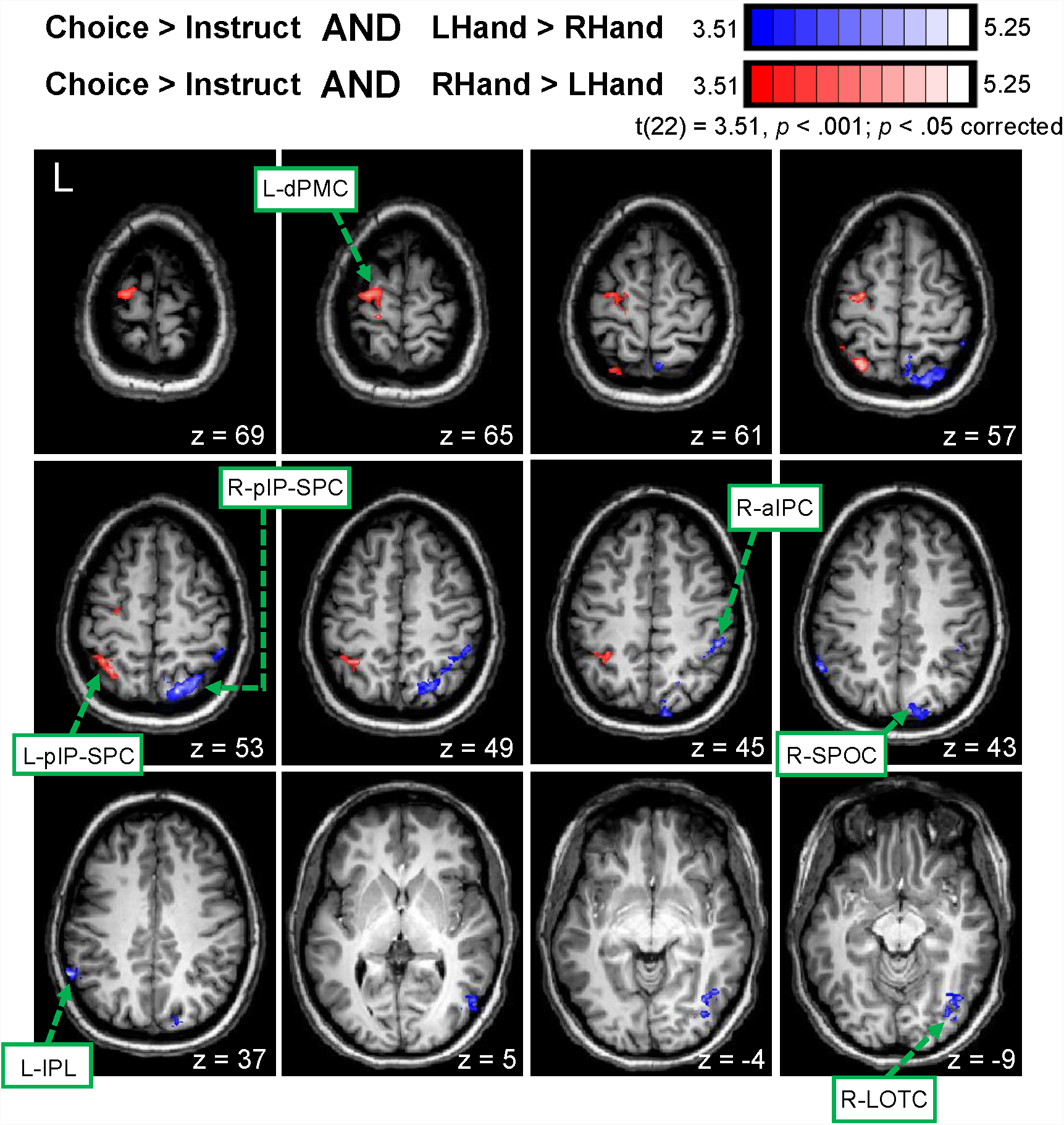
Functional MRI conjunction contrast results: Voxel-wise maps. Statistical activation maps showing significant responses for Choice > Instruct AND LHand > RHand (blue-to-white), and for the complementary conjunction contrast, Choice > Instruct AND RHand > LHand (red-to-white). Group data are shown on the anatomy of a single subject. Brain areas: left dorsal premotor cortex (L-dPMC); left posterior intraparietal and superior parietal cortex (L-pIP-SPC); right posterior intraparietal and superior parietal cortex (R-pIP-SPC); right anterior intraparietal cortex (R-aIPC); right superior parieto-occipital cortex (R-SPOC); left inferior parietal lobule (L-IPL); right lateral occipitotemporal cortex (R-LOTC).

**Figure 5.**
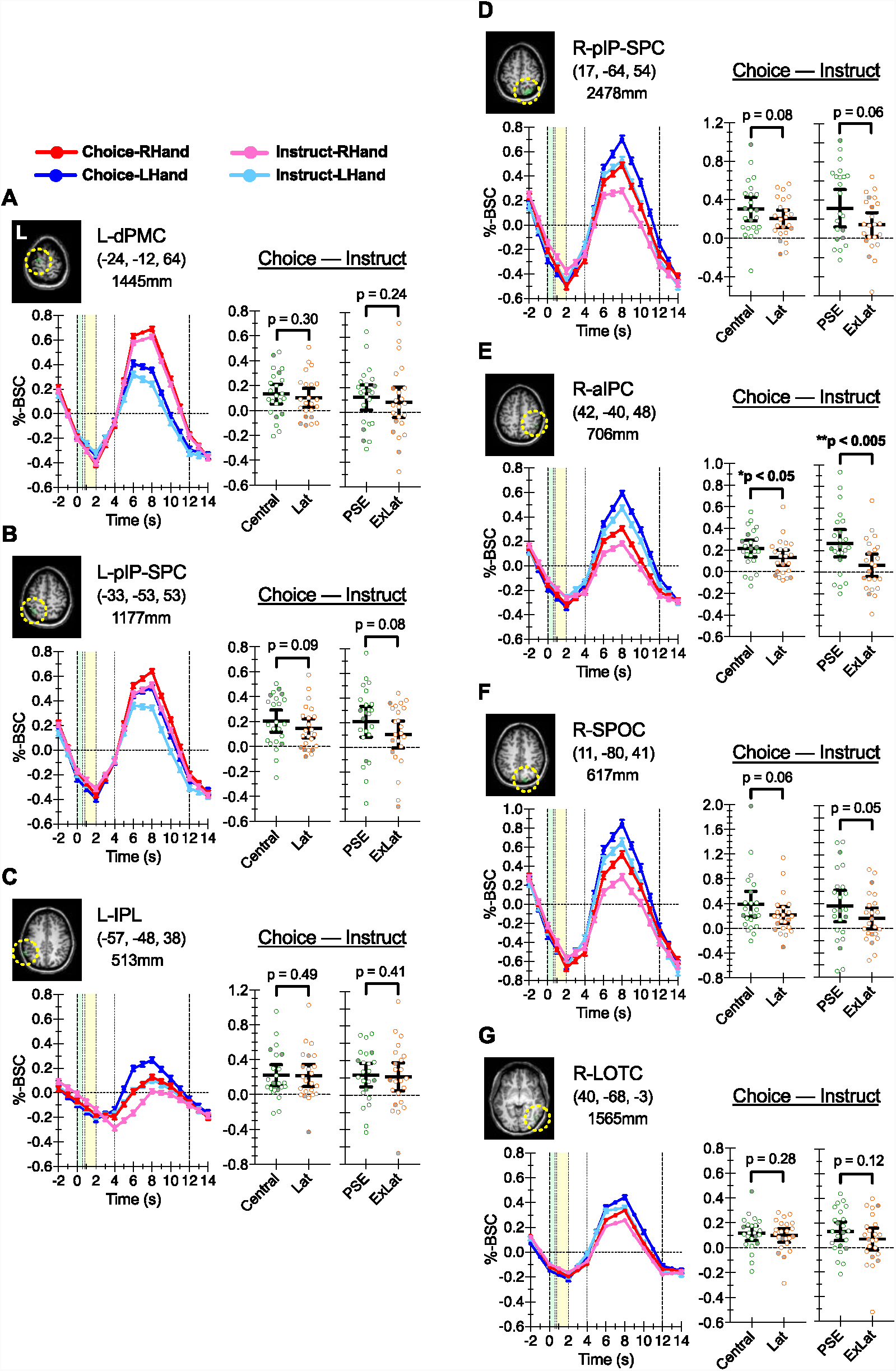
Functional MRI conjunction contrast results: ROI analyses. **(A-G)** Data extracted from areas identified via voxel-wise conjunction contrasts, as reported in Figure 4. Per area, time course data illustrate event-related averaged percent BOLD signal change (%-BSC) values per condition over time, aligned to the onset of the task instruction cue (green shading). The target illumination period is shown in yellow shading. Error bars in the time course data indicate SEMs. Scatter plots indicate %-BSC values expressed as difference scores between Choice – Instruct conditions as a function of Target Location: Central (green) versus Lateral (Lat) (orange); PSE (green) versus Extreme Lateral (ExLat) (orange). Open circles show individual participant scores. Participants without RT data are indicated as filled circles. Solid lines indicate group means with 95% confidence intervals. Brain area abbreviations are defined in Figure 4 caption.

**Figure 6.**
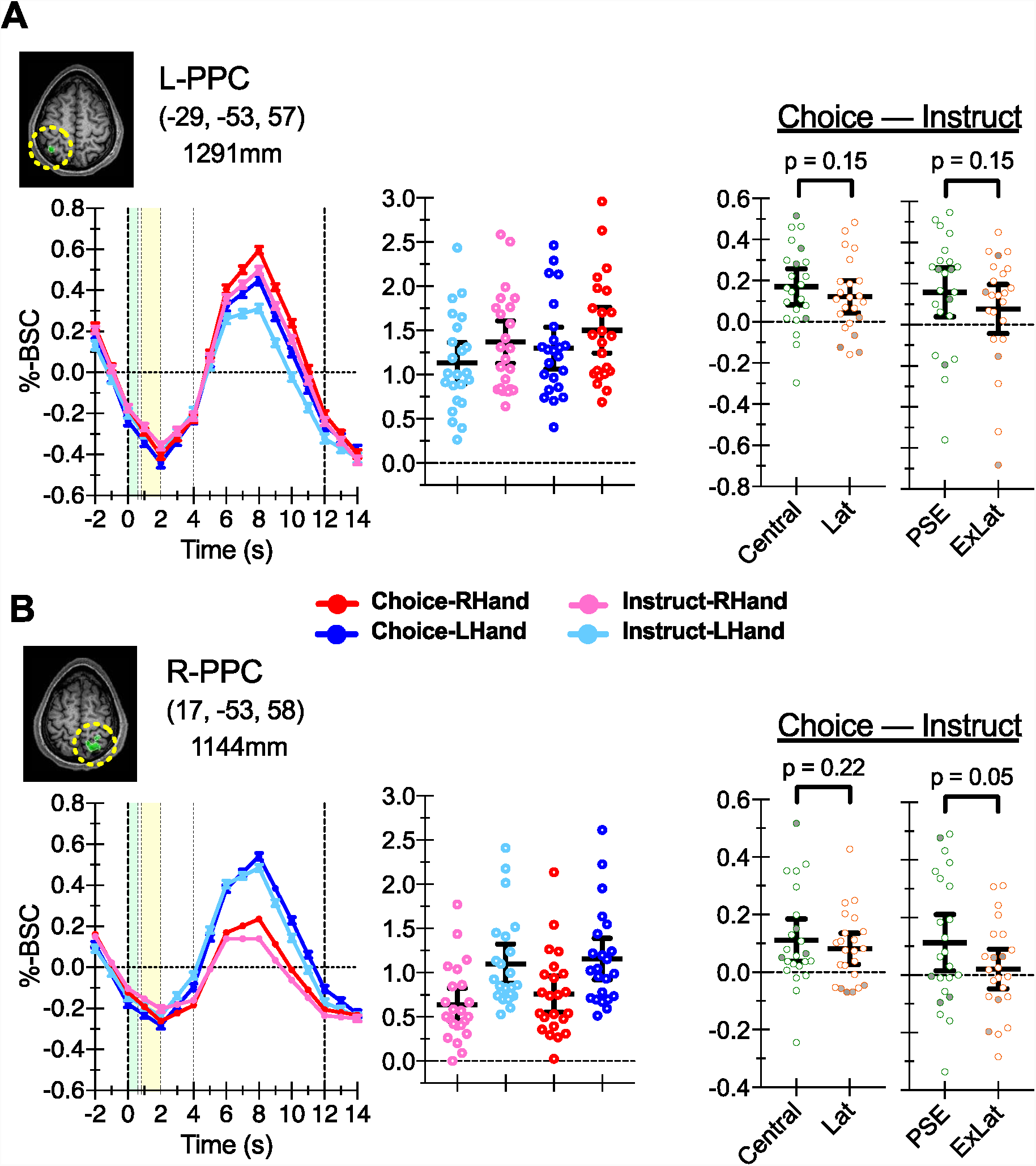
Functional MRI independent ROI results. **(A/B)** Functionally defined L- and R-PPC ROIs, respectively, independently defined on the basis of previous results from Valyear and Frey (2015). Time course data per ROI illustrate event-related averaged percent BOLD signal change (%-BSC) values per condition over time, aligned to the onset of the task instruction cue (green shading). The target illumination period is shown in yellow shading. Error bars in the time course data indicate SEMs. Scatter plots indicate %-BSC values per condition, with individual participant data shown as open circles. Solid lines indicate group means with 95% confidence intervals. The two leftmost scatter plots show %-BSC data expressed as difference scores between Choice – Instruct conditions as a function of Target Location: Central (green) versus Lateral (Lat) (orange); PSE (green) versus Extreme Lateral (ExLat) (orange). Participants without RT data are indicated as filled circles.

Each run comprised 37 trials: 12 Choice, 12 Instruct Left Hand (Instruct-LHand), 12 Instruct Right Hand (Instruct-RHand), and lasted 7min and 30s (225 volumes). The first (“dummy”) trial of each run was discarded from subsequent analyses, since its trial history could not be controlled. Runs included 6s (3 volumes) of rest to begin. Participants performed between 4 to 8 runs (mean = 6.87 runs; mode = 8 runs).

A custom Matlab (R2013b) script was used to create eight distinct run orders where trial history was balanced for each condition within runs. Specifically, 12 targets were presented per condition per run, balanced across Lateral and Central space, with an equal number of targets presented per hemispace, and the order of the presentation of each target position balanced across conditions. The presentation of run orders was pseudo-randomized across participants.

### 2.4. Pre-scan testing

Prior to scanning (mean = 5 ± 7 days, range = 1 to 27), participants took part in a behavioural training session. Training was performed in a mock scanner designed to approximate the same physical constraints as the real MRI scanner but with no magnetic field. The same apparatus and materials used in the real MRI scanner were used for pre-scan testing (Figure 2A). Participants completed a minimum of three, and maximum of four runs. A motion capture system, MoTrak (Psychology Software Tools Inc., 2012; version 1.0.3.4), was used to monitor participant head position during pre-scan testing.

The purpose of the pre-scan testing session was twofold. First, participants learned how to perform the task while keeping their head still. The problems associated with in-scanner head motion were thoroughly explained. Participants were told that their hand actions should involve minimal movements of the upper arm or shoulder, and that their head should be kept still at all times. Actions were trained to be performed smoothly. It is worth emphasizing here that the primary purpose of pre-scan training was to verify that participants could *keep their head still* while performing the task. Otherwise, the task was not difficult to learn or perform. For these reasons, we were unconcerned about large between-subject differences in timing between pre-scan and MRI testing.

Second, pre-scan testing was used to identify and exclude participants who either (1) moved their head too much, or (2) showed little variation in hand choice behaviour. Specifically, participants who showed evidence of excessive/abrupt head movements during the task, or who demonstrated > 75% use of the same hand during the Choice condition did not participate in fMRI testing. We recognize that these procedures introduce selection bias, and that this represents a limitation of our study. However, in the absence of sufficient variation in hand choice behaviour, we would be unable to test our current hypotheses.

Five participants (out of 34) were identified as showing > 75% use of the same hand during the Choice condition, and thus were excluded from fMRI testing. An additional five participants who completed pre-scan behavioural testing were later found to have (safety-related) contraindications for MRI testing, and were excluded.

### 2.5. Imaging parameters

Imaging was performed on a 3-Tesla Philips Achieva MRI scanner with a conventional 8-channel birdcage (SENSE) head coil. Functional MRI volumes were collected using a T2*-weighted single-shot gradient-echo echo-planar imaging (EPI) acquisition sequence: time to repetition (TR) = 2000ms; time to echo (TE) = 30ms; flip angle = 77°; matrix size = 64 by 64; field of view (FOV) = 256mm; slice thickness = 4mm; in-plane resolution = 4mm by 4mm; acceleration factor (integrated parallel acquisition technologies, iPAT) = 2 with parallel acquisition (SENSE). Each volume comprises 38 axial-oblique slices (0.1mm gap), spanning from the most superior point of cortex ventrally to include the entire cerebellum (i.e. whole-brain coverage). A T1-weighted anatomical image was collected using a multiplanar rapidly acquired gradient echo (MP-RAGE) pulse sequence: time to repetition (TR) = 1500ms; time to echo (TE) = 3.45ms; flip angle = 8°; matrix size = 224 by 224; field of view (FOV) = 224mm; 175 contiguous transverse slices; slice thickness = 1mm; in-plane resolution = 1mm by 1mm.

### 2.6. Functional MRI data preprocessing

Imaging data were preprocessed and analysed using Brain Voyager QX (BVQX) version 2.4.2.2070, 64-bit (Brain Innovation, Maastricht, The Netherlands). Each functional run was assessed for subject head motion by viewing cineloop animations and by examining Brain Voyager motion-detection parameter plots after running 3D motion correction algorithms on the untransformed two-dimensional data using BVQX trilinear (motion detection) and sinc interpolation (motion correction) options.

Functional data were preprocessed with linear trend removal and high-pass temporal frequency filtering to remove frequencies below three cycles per run. Functional data were aligned to anatomical volumes, and transformed to standard stereotaxic space (Talairach and Tournoux, 1988). Data were spatially smoothed for group analyses using a Gaussian kernel of 8mm (2 voxels) (full-width at half-maximum).

### 2.7. Data analysis

#### 2.7.1. Matched Choice and Instruct conditions

In-scanner videos were used to match participant’s motor responses between Choice and Instruct conditions. Specifically, for each target position presented within a given run, the hand used to respond during the Choice condition determined which of the two Instruct conditions – LHand/RHand – were defined as ‘matched’, and used for subsequent behavioural and fMRI analyses. For example, if target position 1 (see Figure 2A) involved a left-hand response during the Choice condition, the corresponding Instruct-LHand trial for target position 1 was ‘held’ for analyses – defined as ‘matched’ –, while the Instruct-RHand trial for target position 1 from this same run was excluded from further analyses. This was an essential feature of our design. With this approach, comparisons between Choice and Instruct conditions, for both fMRI and RT data, are equated for motor and visual properties.

Videos were monitored and scored by an experimenter online, and independently scored by two additional experimenters, offline. Specifically, each rater observed participant performance and categorized the following errors: (1) Instruct trials were initiated with the incorrect hand; (2) movements changed abruptly during reaching; (3) no response was made. Errors in performing the task were scored (see Supplementary Table 1), and these trials were excluded from RT analyses, and assigned a predictor of no-interest for fMRI analyses. Rater 1 scored all video data, while Raters 2 and 3 scored video data for a total of 10 and 16 participants, respectively.

**Table 1.** ROI results for areas defined by the voxel-wise conjunction contrasts.

| Brain Area | Hand Specificity<br>(unmatched-Contralateral<br>> unmatched-Ipsilateral) |  | Task by Target<br>Location:<br>Central/Lateral<br>Interaction Term<br>(1, 22) |  | Task by Target<br>Location:<br>PSE/ExLat<br>Interaction Term<br>(1, 22) |  |
| --- | --- | --- | --- | --- | --- | --- |
|  | *t* | *p* | *F* | *p* | *F* | *p* |
| **L-dPMC** | 8.90 | <.001 | 0.53 | 0.24 | 0.53 | 0.24 |
| **L-IPL** | -3.71 | 0.001 | <.001 | 0.49 | 0.05 | 0.41 |
| **L-pIP-SPC** | 3.78 | 0.001 | 1.84 | 0.09 | 2.167 | 0.08 |
| **R-aIPC** | 3.44 | 0.002 | 4.55 | 0.02 | 8.73 | 0.004 |
| **R-LOTc** | 4.44 | <.001 | 0.34 | 0.28 | 1.41 | 0.12 |
| **R-pIP-SPC** | 3.68 | 0.001 | 2.17 | 0.08 | 2.72 | 0.06 |
| **R-SPOC** | 4.28 | <.001 | 2.55 | 0.06 | 2.80 | 0.05 |

#### 2.7.2. Behavioural data analysis

##### Hand choice: Point of subjective equality (PSE)

Hand choice was coded online by an experimenter, and confirmed offline with video and button-release data. To quantify hand choice behaviour per participant, and at the group-level, target locations were reduced from 16 to 8 positions, depending on the lateral distance from midline (Figure 3C), and the point in target space where the use of either hand was equally likely was defined – the Point of Subjective Equality (PSE) (mean number of trials per target per participant = 20.5 trials, ± 0.91 SEM). Specifically, a psychometric function (McKee et al., 1985) was computed according to each participant’s hand choice behaviour per target location, and the PSE was estimated by fitting a general linear model (as described in Valyear et al., 2018). The model contains target positions and a constant term, and uses a Logit link function to estimate the binomial distribution of hand choice responses (1 = right | 0 = left). Model coefficients are evaluated at 1000 linearly spaced points between the outermost values of the target array (i.e. ± 7.6 cm), and the value closest to a 0.50 probability estimate is defined as the PSE.

Pearson’s *r* correlation analysis was used to test for a relationship between PSE and Waterloo handedness scores. A significant negative relationship was hypothesized. Positive Waterloo scores (max = +30) reflect (self-report) right-hand preferences, while negative PSE scores reflect right-hand choice preferences. Outliers were defined as ± 2.5 standard deviations from the group mean and removed from further analysis. Given the directional predictions of this test, we considered a one-tailed p < 0.05 as significant.

##### Response times

Response times (RTs) were defined as the time from the onset of target illumination to the release of (left/right hand) start buttons (i.e. times-to-movement onset). Data from pre-scan training trials were not included in these analyses.

We tested the effects of task instruction, hand used, and target location on RT with linear mixed-effects implemented using the lme4 package (Bates et al., 2014) for R (R Core Team, 2018). Statistical significance was tested for fixed effects by fitting the model with restricted maximum likelihood (REML), deriving degrees of freedom via Satterthwaite approximation using the lmerTest package (Kuznetsova et al., 2017). This approach has shown acceptable levels of Type I error for smaller datasets (<60 items; Luke, 2017). We contrasted levels of significant fixed effects with Tukey adjustment using the lsmeans package (Lenth, 2016).

We tested two models. Each model included the fixed effects of Task (Choice, Instruct) and Hand (LHand, RHand), but differed in how Target Location was defined. In the first model, Target Location was defined as Central (targets 5-12) and Lateral (targets 1-4 and 13-16) conditions. We refer to this model as RT-Central.

The second model was used to test for effects of Target Location defined according to individual-level PSE data. Specifically, Target Location was defined per individual as those targets nearest to the PSE, versus those in the far “extreme” lateral positions (ExLat; targets 1, 4, 13, 16) of the target display, corresponding with ±7.6 cm distances from the midline of the display (Figure 3A). We refer to this second model as RT-PSE.

Both models permitted all possible interactions between fixed effects, and included a random intercept and slope for all fixed effects per subject and a random intercept per run.

We also analysed RT data using repeated measures analysis of variance (RM-ANOVA), and report these data in Supplementary Materials.

#### 2.7.3. Functional MRI data analysis

Analyses were based on a group-level random-effects (RFX) GLM with five predictors specified: Choice-LHand, Choice-RHand, Instruct-LHand, Instruct-RHand, and a predictor of no-interest (i.e. including the first trial of each run, unmatched Instruct trials, and errors). Predictors were modelled as two-volume (four second) boxcar functions aligned to the onset of each trial, convolved with the BVQX default two-gamma function designed to estimate the spatiotemporal characteristics of the Blood-Oxygen-Level Dependent (BOLD) response. Each run was percent-transformed prior to GLM analysis.

A group-level inclusion mask was defined, and used to constrain all subsequent tests. The mask comprised those voxels that were significantly identified by any of the following contrasts: (1) Choice-LHand > rest; (2) Choice-RHand > rest; (3) Instruct-LHand > rest; (4) Instruct-RHand > rest. The resultant statistical activation map was thresholded at *t*(23) = 3.80, p < 0.01 uncorrected, p < 0.05 cluster-size corrected (see Supplementary Figure S1). The purpose of this method was to increase the sensitivity of subsequent statistical tests by reducing the number of voxels considered for correction for multiple comparisons to those that show task-related fMRI activity increases.

##### Voxel-wise conjunction contrasts

The PPIC model specifically predicts a main effect of Task (Choice > Instruct) and a main effect of Hand (Contralateral > Ipsilateral) within bilateral pIP-SPC (Figure 2B). We use the following two conjunction contrasts to directly test these predictions:

(1).(Choice-LHand + Choice-RHand) > (Instruct-LHand + Instruct-RHand) AND (Choice-LHand + Instruct-LHand) > (Choice-RHand + Instruct-RHand)

This conjunction tests for areas showing Choice > Instruct and LHand > RHand, predicted to identify the right hemisphere pIP-SPC (R-pIP-SPC).

(2).(Choice-LHand + Choice-RHand) > (Instruct-LHand + Instruct-RHand) AND (Choice-RHand + Instruct-RHand) > (Choice-LHand + Instruct-LHand)

This conjunction tests for areas showing Choice > Instruct and RHand > LHand, predicted to identify the left hemisphere pIP-SPC (L-pIP-SPC).

Resultant activation maps were set to a statistical threshold of *t* = 3.51 (p < 0.005, one-tailed), corrected for multiple comparisons using Brain Voyager QX cluster-level statistical threshold estimator, found to indicate a minimum cluster size of (1) 298 mm^3^ and (2) 325 mm^3^ (p < 0.05) for each conjunction contrast defined above, respectively.

#### Region-of-interest (ROI) analyses

Multiple ROI-based analyses were performed. In all cases, mean percent BOLD signal change (%-BSC) values, represented as beta weights per condition of interest were extracted from each ROI, and tested. Hand specificity tests (Results section 3.2.2) involved extraction of beta weights corresponding with unmatched Instruct trials from ROIs identified by voxel-wise conjunction contrasts, and comparisons between unmatched-LHand versus unmatched-RHand conditions using paired-samples t-tests, with p < 0.05 taken as significant. These data are independent of the data used to define ROIs.

Task by Target Location ROI-based analyses (Results section 3.2.3) involved testing the RM-ANOVA interaction terms according to our a priori directional hypothesis: (1) (Choice-Central > Instruct-Central) > (Choice-Lateral > Instruct-Lateral); (2) (Choice-PSE > Instruct-PSE) > (Choice-ExLat > Instruct-ExLat). Here, we considered a one-tailed p < 0.05 as significant, given our predictions. These tests are orthogonal to the contrasts used to define ROIs.

Finally, we performed additional ROI analyses on the basis of our prior data showing fMRI repetition suppression for repeated hand actions within bilateral posterior parietal cortex (Valyear and Frey, 2015). Mean %-BSC values from the current data set were extracted from the complete set of active voxels identified from Valyear and Frey (2015) – comprising the ROIs: L-PPC, and R-PPC (Figure 6). This prior investigation involved an entirely different group of participants, and thus, these ROIs were defined independently from the current data.

## 3. Results

### 3.1. Behavioural results

Video data confirm that the task was performed correctly, and reveal very few errors (Supplementary Table 1). Button release data is unavailable for four participants, due to technical errors.

#### 3.1.1. Hand choice

Participants use both hands to respond to targets during the Choice condition, and there is a clear relationship between Hand and Target Position. Expressed as a function of quadrants of the target display (Figure 3B) – left-Lateral (targets 1-4), left-Central (targets 5-8), right-Central (targets 9-12), right-Lateral (targets 13-16) –, the group data reveal that the left hand is typically used for targets in the left-Lateral quadrant, and the right hand is typically used for targets in the right-Central and right-Lateral quadrants, to the right of midline (Figure 3B). Responses to the left-Central quadrant tend to involve a mixture of left- and right-hand responses. These differences were verified via a RM-ANOVA of arcsine transformed proportions of right-hand use (see Supplementary Materials).

Subsequent analyses redefine target space as 8 conditions representing lateral distances from the midline, and reveal a group mean PSE – where the probability of hand choice is balanced between hands – of −1.30 cm, reflecting a leftward (right-hand) bias (Figure 3C). The spread of individual-participant-PSE values includes −6.23 to 0.65, and for the majority of participants, overlaps with left- and right-central quadrants.

Correlation analyses between PSE and Waterloo handedness scores reveal a significant negative relationship (*r* = −0.40, p < 0.05). These results suggest that the leftward shift in PSE reflects the influence of hand preference – as a group, individuals are more likely to choose their preferred (right) hand to reach to targets.

#### 3.2.1. Response times: Linear mixed-effects models

RT data are based on N = 19 participants.

The RT-Central model is a significantly better fit than a null model containing only its random effects (χ^2^ = 76.0, p < 0.001), and reports a significant influence of Task (F(1, 19.9) = 112.9, p < 0.001). RTs are greater for the Choice versus Instruct condition (Figure 3D/E/F), consistent with the additional time required to decide which hand to use – i.e. significant choice costs.

Two additional significant results are revealed. First, RTs are affected by an interaction between Task and Hand (F(1, 3108) = 17.0, p < 0.001). This reflects greater choice costs (Choice > Instruct) for the LHand, although choice costs are significant for both hands (Figure 3D). Specifically, compared with the RHand, RTs are smaller with the LHand for the Instruct condition, yet larger with the LHand for the Choice condition.

Second, RTs are affected by an interaction between Hand and Target Location (F(1, 3104) = 12.1, p < 0.001). This result reflects a non-significant positive difference between LHand-Central – LHand-Lateral (p = 0.13) combined with a non-significant negative difference between RHand-Central – RHand-Lateral (p = 0.88). It is difficult to interpret these results, since the pairwise comparisons are both non-significant. No other significant effects are identified.

Contrary to our predictions, the interaction between Task and Target Location is non-significant (F(1, 3117) = 0.154 p = 0.695) (Figure 3E). These results indicate that the choice costs (Choice > Instruct) are similar for reaches to Central and Lateral targets.

We tested a second model – the RT-PSE model – instead defining Target Location per individual as those targets nearest to the PSE, versus ExLat (targets 1, 4, 13, 16; ±7.6 cm from the display midline). This model was also a significantly better fit for RTs than a null model omitting the fixed effects (χ^2^ = 52.5, p < 0.001). Consistent with the results for RT-Central model, described above, these analyses indicate that RTs are significantly influenced by Task (F(1, 20.8) = 101.2, p < 0.001), and by an interaction between Task and Hand (F(1, 1508) = 8.04, p < 0.001).

The results of the RT-PSE model also reveal a non-significant trend for the interaction between Task and Target Location F(1, 1508) = 2.80, p = 0.09) in the predicted direction: (Choice-PSE > Instruct-PSE) > (Choice-ExLat > Instruct-ExLat) (Figure 3F). Although not passing statistical significance, these results are consistent with the PPIC model, and other bounded-accumulation models (Cisek, 2006; Beck et al., 2008; Hanks et al., 2015), and are interpreted as evidence for a gradient of high (PSE) versus low (ExLat) areas of competition as a function of Target Location. No other significant effects are identified.

### 3.2. Functional MRI results

Participants were able to perform the task in the MRI scanner while keeping their head still (see Supplementary Figure S2 for complete details).

#### 3.2.1. Voxel-wise conjunction contrasts

The PPIC model predicts that bilateral pIP-SPC will respond preferentially to the Choice (> Instruct) and Contralateral (> Ipsilateral) conditions. Consistent with these predictions, the conjunction contrast Choice > Instruct AND LHand > RHand identifies significant activity within the right posterior intraparietal and superior parietal cortex (R-pIP-SPC), while the complementary conjunction contrast, Choice > Instruct AND RHand > LHand identifies significant activity within the left posterior intraparietal and superior parietal cortex (L-pIP-SPC) (Figure 4). Activity within the right hemisphere extends along the intraparietal sulcus, and includes distinct foci within the anterior intraparietal cortex (R-aIPC) and the superior parieto-occipital cortex (R-SPOC), medially, just anterior to the parieto-occipital sulcus. Activity within the L-pIP-SPC is comparatively more focal, largely restricted to intraparietal cortex.

The conjunction contrasts identify three additional brain areas (Figure 4). First, the contrast Choice > Instruct AND RHand > LHand reveals significant activity within the left dorsal premotor cortex (L-dPMC), at the junction of the precentral and superior frontal sulci. Second, the complementary conjunction contrast Choice > Instruct AND LHand > RHand identifies significant activity in two other areas: right lateral occipitotemporal cortex (R-LOTC), overlapping with the posterior middle temporal gyrus, dorsally, and the fusiform cortex, ventrally; left inferior parietal lobule (L-IPL), at the intersection of the supramarginal and angular gyri. The L-IPL is the only area identified that shows stronger activity for responses made with the ipsilateral hand.

The event-related averaged %-BSC time-courses verify the timing of the effects within each area identified by the conjunction contrasts (Figure 5). This step is important to rule out possible differences between conditions that may arise prior to trial onsets; for example, related to previous trial history.

#### 3.2.2. ROI results: Task by Target Location

A priori, we predicted that responses to Central versus Lateral targets would represent more ambiguous hand-use choices by virtue of the greater degree of inter-manual similarity in biomechanical and energetic costs associated with reaching to these target locations – relatively low bias, high competition (Figure 2C). This difference would drive greater fMRI-activity-level differences between Choice and Instruct conditions in bilateral pIP-SPC.

Our fMRI data support these predictions. The patterns of %-BSC values extracted from four areas: L-and R-pIP-SPC, R-aIPC and R-SPOC are consistent with the predicted Task by Target Location interaction – i.e. (Choice-Central > Instruct-Lat) > (Choice-Lateral > Instruct-Lat) (Figure 5; Table 1). These effects reach statistical significance in R-aIPC, and near significance in areas R-SPOC (p = 0.06), L-pIP-SPC (p = 0.09) and R-pIP-SPC (p = 0.08). These results dissociate from our RT data, described above, where no statistical differences in choice-costs (Choice > Instruct) between Central and Lateral Target Locations are identified.

When Target Location is defined per individual as those nearest to the PSE versus ExLat positions, similar findings are obtained. Again, fMRI response levels in bilateral pIP-SPC, R-aIPC and R-SPOC show the predicted Task by Target Location interaction: (Choice-PSE > Instruct-PSE) > (Choice-ExLat > Instruct-ExLat) (Figure 5; Table 1). This pattern of responses is specific to these brain areas, and is consistent with a competitive process underlying hand choice. Choice-costs are higher for responses made to targets near the PSE, where there is minimal bias in hand choice behaviour, and this is associated with significantly more pronounced differences in fMRI response levels between Choice and Instruct conditions. These fMRI data parallel our RT data, showing prolonged RTs for reaches to targets near the PSE for the Choice but not Instruct conditions (although as reported above, the RT data do not reach statistical significance; p = 0.09).

It is important to recognize that our tests involving the PSE versus ExLat conditions were unplanned, and in the case of our fMRI data, may be insufficiently powered; our experimental design provides limited numbers of trials per Task per Lateral Target Location per run. Low numbers of trials per condition per run is problematic for fMRI analyses. Given these limitations, these data should be interpreted cautiously. It is also possible, however, that these experimental-design limitations contribute to the relatively weak statistical significance of these effects.

#### 3.2.3. ROI results: Hand specificity

Our voxel-wise conjunction contrasts identify areas showing both Choice > Instruct and Contralateral > Ipsilateral specificity (aside from the L-IPL, which shows stronger responses for actions with the ipsilateral hand). However, since hand-use and target location are tightly associated – i.e. the majority of left hand reaches are to targets in left hemispace, while the majority of right hand reaches are to targets in right hemispace – interpretation of the Contralateral > Ipsilateral results is confounded. These effects may reflect specificity for actions/stimuli in contralateral hemispace.

To test this hypothesis, from each ROI identified by our conjunction contrasts we extract data representing unmatched Instruct trials, and compare unmatched-LHand versus unmatched-RHand conditions. Critically, these data are independent from those used to define the ROIs. The results reveal significantly greater fMRI responses for the use of the Contra-versus Ipsilateral hand, for all areas identified (aside from the L-IPL, which shows significantly greater fMRI responses for the use of the Ipsilateral – left – hand) (Table 1). Together with the conjunction contrast results, our data demonstrate hand specificity in these brain areas, independent of the spatial locations of targets in the display.

#### 3.2.4. ROI results: Independent tests of the PPIC model

Previous fMRI results from our lab (Valyear and Frey, 2015) constrain the anatomical specificity of the PPIC model to the posterior intraparietal and superior parietal cortex, bilaterally, and motivate two additional functional constraints: (1) hand-specific encoding, and (2) graded contralateral specificity. In other words, our model draws explicitly from these previous data; these same brain areas identified within bilateral posterior parietal cortex – labelled here as L- and R-PPC – are predicted to show both Choice > Instruct and Contralateral > Ipsilateral responses.

To test these predictions, we extract the mean %-BSC values corresponding with Choice-LHand, Choice-RHand, Instruct-LHand, Instruct-RHand conditions from the complete set of active voxels identified within the L- and R-PPC on the basis of our previous study (Valyear and Frey, 2015), and enter these data into a Task by Hand RM-ANOVA. As predicted by the PPIC model, the results reveal significantly stronger responses for both the Choice (> Instruct) and the Contralateral (> Ipsilateral) conditions within both the L- and R-PPC (Figure 6; Table 2).

**Table 2.** ROI results for areas independently defined on the basis of previous fMRI data (Valyear and Frey, 2015).

| ROIs defined by<br>Valyear and Frey<br>(2015) | Hand by Task |  |  |  |  |  | Task by Target<br>Location:<br>Central/Lateral |  | Task by Target<br>Location:<br>PSE/ExLat |  |
| --- | --- | --- | --- | --- | --- | --- | --- | --- | --- | --- |
|  | ME Task<br>(1, 22) |  | ME Hand<br>(1, 22) |  | Interaction Term<br>(1, 22) |  | Interaction Term<br>(1, 22) |  | Interaction Term<br>(1, 22) |  |
| Brain Area | F | p | F | p | F | p | F | p | F | p |
| **LH-PPC** | 19.31 | <.001 | 25.49 | <.001 | 1.14 | 0.30 | 1.10 | 0.15 | 1.10 | 0.15 |
| **RH-PPC** | 11.32 | 0.003 | 126.73 | <.001 | 4.70 | 0.04 | 0.60 | 0.22 | 2.91 | 0.05 |

Post-hoc comparisons confirm greater responses for the Choice versus Instruct conditions for both Contra- and Ipsilateral conditions, within both L- and R- PPC (Figure 6). This is an important aspect of our findings, consistent with the PPIC model and the hypothesis that action plans for both hands are represented bilaterally within pIP-SPC.

We also test for effects of Task by Lateral Target Location, according to both Central versus Lateral, and PSE versus ExLat conditions, respectively. The trends in both ROIs, though non-significant, are in the predicted directions, and in particular, reach near statistical significance (p = 0.05) in the R-PPC for the predicted (interaction) pattern of (Choice-PSE > Instruct-PSE) > (Choice-ExLat > Instruct-ExLat) (Figure 6; Table 2).

## 4. Discussion

The current data significantly advance our understanding of human hand choice behaviour. Few previous studies have investigated the brain mechanisms involved in ‘free choice’, and instead involve action selection on the basis of arbitrary rules. This is the first brain imaging study to investigate free hand choice in humans. Our findings reveal the selective involvement of a network of brain areas within bilateral posterior parietal cortex, left-lateralized inferior parietal and dorsal premotor cortices, and right lateral occipitotemporal cortex.

At the outset, we formulate a systems-level model of hand choice, the Posterior Parietal Interhemispheric Competition (PPIC) model. The model generates specific predictions, and provides a useful conceptual framework to constrain our results interpretations. We first evaluate our data within this framework, and then interpret the significance of our results revealing hand-choice selectivity in additional brain areas, not predicted by the model.

### The PPIC model

According to the Affordance Competition Hypothesis (Cisek, 2007), the neural mechanisms that specify action possibilities in sensorimotor terms also play an important role in selecting among those possibilities. Areas within monkey superior parietal (Caminiti et al., 1996; Scherberger et al., 2005) and dorsal premotor (Scott et al., 1997; Hoshi and Tanji, 2004) cortices are necessary for the transformation of visual information to motor commands for reaching, and critically, the neural responses within these areas also reflect reach choices (Cisek and Kalaska, 2005; Scherberger and Andersen, 2007; Pesaran et al., 2008; Pastor-Bernier and Cisek, 2011; Thura and Cisek, 2014). Temporary inactivation of the “parietal reach region” – area PRR, located within the medial bank of the intraparietal sulcus – impairs reach (but not saccade) selection (Christopoulos et al., 2015b). These data provide powerful evidence for the causal involvement of the PPC in reach choices.

The PPIC model borrows from the neural population-level response dynamics specified by the Affordance Competition Hypothesis, and extends these principles to hand-specific encoding and hand selection. Neural populations within bilateral pIP-SPC encode possible actions in hand-specific terms and compete for selection across and within hemispheres. Actions with either hand are represented bilaterally, yet within each hemisphere the contralateral hand is overrepresented.

Consistent with the PPIC model, our findings reveal the involvement of bilateral pIP-SPC in hand choice. Responses within pIP-SPC are significantly greater for the Choice versus Instruct condition, when hand use is freely selected. These effects are not attributable to motor or visual confounds, including potential differences in motor- or visual-response sensitivity to targets presented at different spatial locations. Choice and Instruct conditions are carefully matched for responses to each target location so that the contrast between these conditions is balanced for these features.

These same brain areas demonstrate a pattern of graded contralateral response specificity. Responses are strongest for actions made with the contralateral hand; although, actions with the ipsilateral hand also yield robust responses. Further, differences between Choice and Instruct conditions are not restricted to responses made with the contralateral hand. The Choice condition preferentially activates bilateral pIP-SPC, even for ipsilateral responses. This pattern is consistent with a role for the planning and selection of actions with either hand, as specified by the PPIC model.

The anatomical specificity of our data is consistent with the PPIC model, and the hypothesis that hand selection involves the same brain areas that are important for action planning. Bilateral pIP-SPC and R-SPOC showing preferential responses for the Choice condition closely overlap with areas implicated in the planning and sensorimotor control of the arm for reaching (Astafiev et al., 2003; Connolly et al., 2003; Medendorp et al., 2005; Prado et al., 2005; Culham and Valyear, 2006; Tosoni et al., 2008; Fabbri et al., 2010; Pitzalis et al., 2010; Vesia and Crawford, 2012; Andersen et al., 2014; Monaco et al., 2015). Consistent with our data, Beurze et al. (2007) demonstrate that during the planning phase of a reaching task, bilateral pIP-SPC integrates information about the spatial location of targets with the hand that will be used for reaching.

Finally, our results provide evidence for a competitive process underlying hand choice. Responses in bilateral pIP-SPC demonstrate increased levels of choice-specificity (Choice > Instruct) for reaches made to targets near the midline (Central) compared to the left/right (Lateral) edges of the display. These data are consistent with a gradient of increased levels of competition between neural populations representing hand-specific reach plans for targets near the midline, where inter-manual differences in the biomechanical and energetic costs associated with reaching are minimal.

Unexpectedly, however, our behavioural RT data reveal a more complex relationship between choice-costs and target location. Although RTs indicate significant choice-costs (Choice > Instruct), these costs are similar for reaches to Central and Lateral targets. Additional analyses indicate that for most participants the area in target space of maximal hand-choice ambiguity is shifted to the left of midline. This represents the theoretical point in target space where the use of either hand was equally probable – the PSE –, and a significant correlation between participant PSE and Waterloo handedness-preference scores suggests that this leftward shift reflects the influence of hand preference. Analyses of RT data indicate a non-significant (p = 0.09) trend in the predicted direction of greater choice-costs – greater Choice > Instruct differences – for reaches to targets near the PSE.

Complementary fMRI analyses reveal response patterns within bilateral pIP-SPC, R-aIPC, and R-SPOC that parallel these RT data – the strength of the Choice > Instruct differences in fMRI response levels in these brain areas are more pronounced for reaches to targets near the PSE. These particular aspects of our results should be interpreted cautiously, however. At this level, we may have too few trials per condition to reliably estimate fMRI responses. Notwithstanding these limitations, our PSE-level analyses reveal congruent fMRI and RT results that are consistent with the PPIC model, and a competitive process underlying hand choice. Choice-costs are higher for reaching to parts of target space where there is minimal bias in hand choice behaviour.

Although speculative, we suggest that our discrepant findings between RT and fMRI data regarding the influence of Central versus Lateral target locations relate to differences in how biomechanical factors interact with hand preference to influence these measures. According to the PPIC model, Lateral versus Central target locations represent a narrower range of reach possibilities, and thus will activate fewer competing neural populations encoding those possibilities. As a consequence, the number of active neural units in competition, the time required for the activity of one population to reach suprathreshold levels, and the number of neural units that are actively inhibited after threshold is reached are reduced. All three of these factors will drive down fMRI response levels, while only the second factor – decreased times to reach threshold – will influence RTs. This can explain why, compared with RTs, fMRI data may show pronounced effects of target location.

According to these factors, however, RTs and fMRI activity-levels should nonetheless follow the same direction. Our Central-Lateral data do not. To explain this discrepancy, we suggest that hand preference influences hand choice by driving changes in the accumulation-to-threshold rates of competing neural units, and disproportionately influences RTs compared with fMRI activity levels. For Central targets in our display, increased accumulation-to-threshold rates in neural populations encoding the preferred (right) hand will reduce decision times and lead to the predominate use of the preferred hand. Despite these changes, however, the number of active neural units in competition, and the number of neural units that are actively inhibited after threshold is reached remain high. These differences, at least in principle, could explain why our fMRI data reveal greater Choice > Instruct effects for Central versus Lateral target locations while our RT data do not.

Other data are consistent with the current findings, and support the concept of simultaneously active reach plans competing for selection. When reaching to multiple potential targets, human behavioural (Gallivan et al., 2016; Gallivan et al., 2017), and monkey neurophysiological (Cisek and Kalaska, 2005; Scherberger and Andersen, 2007; Pastor-Bernier and Cisek, 2011) data suggest that parallel action plans are specified in motor (not visual) coordinates, and compete for selection. Further, although these studies tend to investigate reach choices involving the same effector, recent data suggest that similar “action-based” competitive models can explain effector-selection (Christopoulos et al., 2015a). Temporary inactivation of reach-(Christopoulos et al., 2015b) versus saccade-selective (Christopoulos et al., 2018) areas in monkey posterior parietal cortex (areas PRR, mentioned above, and the lateral intraparietal area, LIP, respectively) selectively impairs reach versus saccade choices, respectively, and these data can be explained by a computational model that specifies competitive interactions between these brain areas (Christopoulos et al., 2015a). Conceptually, our PPIC model is consistent with this framework. In the PPIC model, parallel competitive interactions take place between brain areas in the PPC encoding hand-specific action plans, and mediate hand choice.

Our findings complement and extend those of Oliveira et al. (2010). Using single-pulse TMS, Oliveira et al. (2010) demonstrate a necessary role for the left PPC in hand choice. TMS to left PPC during the planning phase of a free-choice reaching task is shown to shift the probability of choices in favour of increased use of the left hand. Conversely, stimulation to the right PPC had no significant influence on hand choice. This asymmetry was unexpected, and the authors offered several possible explanations. Our new findings help to disentangle these different explanations.

First, Oliveira et al. (2010) speculate that perhaps the left-but not the right-hemisphere PPC represents action plans with both hands, and can therefore compensate for the disruptive effects of TMS to right PPC. Our data are inconsistent with this account, however. We find that both the L- and R-pIP-SPC respond preferentially when hand choice is necessary, and for both contra- and ipsilateral responses. If the right hemisphere PPC only represents action plans with the contralateral hand, preferential activity for the Choice condition for the ipsilateral hand is unexpected.

As another possibility, Oliveira et al. (2010) suggest that the critical functional area involved in hand choice may be more spatially restricted within the right PPC, and thus was not effectively disrupted via their TMS manipulation. Our data are inconsistent with this account, also. We find relatively widespread involvement of the right hemisphere pIP-SPC in hand choice. If the critical area in right PPC was ‘missed’ by Oliveira et al. (2010), our data suggest that this was unlikely the consequence of spatially more circumscribed involvement of the right PPC in hand choice.

Finally, Oliveira et al. (2010) recognize that the absence of reliable right PPC TMS effects may relate to the strong right-hand bias present in their group of right-handers tested. This may have left little room for increased use of the right hand, following right PPC stimulation. Although our data do not directly address this possibility, this account remains tenable and represents an important hypothesis for future studies to investigate.

### Visuospatial interpretations

Our data reveal the involvement of bilateral pIP-SPC in hand selection, and demonstrate that these areas show contralateral hand specificity, more robustly activated for actions made with the contralateral hand. Given that in our paradigm hand choice and space are closely associated, however, it is important to consider an account of the contralateral specificity of fMRI responses within bilateral pIP-SPC as attributable to visuospatial rather than (hand-specific) motor coding. Specifically, since reaches with the left hand are predominately made to targets in left hemispace, and vice-versa for right-hand reaches, contralateral specificity within bilateral pIP-SPC may reflect preferential neural responses for targets in contralateral hemispace, rather than the specification of hand-specific action plans.

Critically however, additional analyses controlling for target space confirm significant preferential fMRI responses for actions with the contralateral hand within L- and R-pIP-SPC. These data are not attributable to visuospatial coding, and instead reflect genuine contralateral hand-specificity. Also, preferential fMRI responses for the Choice condition in bilateral pIP-SPC are evident for actions made with the ipsilateral hand, a pattern that conflicts with a strictly visuospatial encoding account, but that is consistent with the PPIC model.

### Additional brain areas

Alongside bilateral pIP-SPC, our results indicate the involvement of left dorsal premotor cortex (L-dPMC), left inferior parietal lobule (L-IPL), and right lateral occipitotemporal cortex (R-LOTC) in hand choice. All areas demonstrate significantly stronger activity for the Choice versus Instruct conditions. L-dPMC and R-LOTC are also more strongly activated for reaching with the contralateral hand, while the L-IPL is more strongly activated for reaching with the ipsilateral hand.

The dPMC is densely interconnected with intraparietal and superior parietal areas, and together these areas mediate the planning and online control of reaching (Scott et al., 1997; Wise et al., 1997; Vesia et al., 2005). The involvement of dPMC in the planning and selection of reaching actions is predicted by the Affordance Competition Hypothesis (Cisek, 2007), and supported by various data (reviewed above). Graded contralateral specificity within dPMC is also consistent with previous data (Medendorp et al., 2005; Beurze et al., 2007). The significance of the left-lateralization of these results is unclear, although previous findings indicate a predominant role for the left hemisphere in action selection (Schluter et al., 2001; Rushworth et al., 2003; Koch et al., 2006; Jacobs et al., 2010).

In the absence of advance predictions about the involvement of the R-LOTC and L-IPL in hand choice, we can only speculate as to the significance of these results. The importance of the LOTC in high-level visual processing is well established (Grill-Spector and Malach, 2004). Our activity in the R-LOTC likely includes the Extrastriate Body Area (EBA), a functionally-defined, predominately right-lateralized region within LOTC that is preferentially responsive to viewing human bodies (versus other object categories) (Downing et al., 2001). Although part of the ventral visual pathway (Ungerleider, 1982; Goodale and Milner, 1992), and considered essential for body-part visual perception and recognition (Urgesi et al., 2004), other data suggest a role for the EBA in action planning. The spatial patterns of fMRI responses within EBA reliably distinguish between different types of upcoming actions performed with the hand (Gallivan et al., 2013), and the EBA is active during the performance of reaching actions in the absence of visual feedback (Astafiev et al., 2004; Orlov et al., 2010). These previous findings suggest that R-LOTC is not only important for high-level visual processing, but also plays a role in action planning. Our data extend this hypothesis to suggest that the R-LOTC is also important for hand choice.

The left supramarginal gyrus has long been associated with limb praxis and the performance of learned actions (Buxbaum, 2001; Goldenberg, 2009), including a specific role for action planning and selection (Buxbaum et al., 2005), while other data also implicate this area as important for visuospatial attention, and in particular, attentional reorienting (Corbetta et al., 2005). Our findings reveal the involvement of the L-IPL in hand choice, and in particular, during free choice actions made with the left hand. Although speculative, the preferential engagement of this area for reach-choices made with the left hand may reflect increased processing demands related to the selection and use of the non-preferred hand. Future studies involving free hand choice with both left- and right-handed participants will be of value.

These aspects of our results motivate changes to our proposed model. Alongside the involvement of bilateral posterior intraparietal cortex, our data indicate that the L-dPMC, L-IPL and R-LOTC are important for deciding which hand to use to perform actions. Further understanding how this network interacts to govern hand choice, and the potentially distinct functional contributions of these different brain areas, is an important goal for future research.

### Concluding remarks

The brain mechanisms involved in ‘free choice’ have been scarcely studied; most previous investigations focus instead on rule-based action selection, where the mappings between stimuli and responses are arbitrary (e.g. respond with the left hand when a stimulus is a particular colour). Here we identify a network of brain areas involved in selecting which hand to use to perform actions on the basis of ‘natural’ factors – e.g. target location –, similar to the conditions that commonly constrain these choices in everyday life. Our data reveal the specific involvement of bilateral posterior intraparietal and superior parietal cortex, left dorsal premotor cortex, left inferior parietal lobule, and the right lateral occipitotemporal cortex. Our findings provide support the PPIC model, and the hypothesis that hand-specific action plans are concurrently activated in bilateral posterior parietal cortex, and compete for selection. We suggest that, although incomplete, the PPIC model of hand choice is of continuing heuristic value, and warrants further investigation.

